# Testosterone alleviates inflammation but increases the methacholine response in mice with allergic lung inflammation

**DOI:** 10.64898/2026.02.10.705120

**Authors:** Cyndi Henry, Magali Boucher, Andrés Rojas-Ruiz, Louane Camillari, Louis Gélinas, Marie-Josée Beaulieu, David Marsolais, Vincent Joseph, Ynuk Bossé

## Abstract

Testosterone seems protective against asthma, but the underlying mechanisms are uncertain. Herein, the effect of testosterone was investigated on several features of experimental asthma. Systemic testosterone was first altered to subphysiological, physiological, or supraphysiological levels in male BALB/c mice through orchiectomy and testosterone supplementation. Testosterone (0.25 mg/day/30 g of body weight) was delivered continuously during 20 days using an implanted pump. At day 10, each group was exposed intranasally to either saline or house dust mite (HDM) once daily for 10 consecutive days to induce allergic lung inflammation. The day after the last exposure, respiratory mechanics was measured at baseline and in response to nebulized methacholine. Bronchoalveolar lavages (BAL) and lung tissues were also collected to quantify inflammation. Baseline respiratory mechanics were altered in mice with subphysiological levels of testosterone, with signs of small airway narrowing heterogeneity and closure. Testosterone drastically inhibited the HDM-induced inflammation. Yet, testosterone also increased the response to methacholine, as well as hysteresis, which are both indicators of enhanced airway smooth muscle activity. While it suggests that testosterone increases the contractility of the smooth muscle, it simultaneously and markedly inhibits inflammation. Explanations as to how these outcomes may lead to protection in asthma are discussed.

## INTRODUCTION

Asthma is more prevalent in males than females during childhood^1-3^. However, a switch occurs during adolescence when asthma becomes more prevalent in females^1-3^. This switch occurs mainly because of a plummeting prevalence in males that coincides with a rise in circulating testosterone levels^4^. This trend is worldwide and, together with positive associations between androgen levels and favorable asthma outcomes^5,6^, it represents indirect evidence that testosterone is protective against asthma.

Several studies have investigated the effect of testosterone in mouse models of asthma. It seems clear that testosterone downregulates allergic lung inflammation^7-17^. Yet, testosterone also increases the response to nebulized methacholine in healthy mice^18,19^, which is consistent with the greater methacholine response seen in males *versus* females^20-24^ and the decreased methacholine response in males post-orchiectomy^18,19,25^.

Collectively, these observations are baffling because asthma and hyperresponsiveness to methacholine are intimately linked^26,27^, implying that factors increasing the methacholine response are expected to contribute to asthma as well. This is clearly not the case for testosterone. While it increases the methacholine response in healthy mice^18,19^, studies in both humans^1-3,5,6^ and mice^7-17^, as well as an early trial with testosterone in humans^28^, suggest that testosterone is protective against asthma.

Clearly, more studies are needed for deciphering the intricacies by which testosterone protects against asthma. In the present study, not only mice with or without experimental asthma were studied after orchiectomy or the sham surgery, but additional groups with orchiectomy were supplemented with a continuous supply of testosterone. This yielded mice with three levels of circulating testosterone during the entire experimental protocol, as well as a clear edge for evaluating how the magnitude and directionality of its effect on inflammation, respiratory mechanics, and the methacholine response play out to protect against experimental asthma.

## METHODS

### Mice

Ninety-six male BALB/c mice (Charles River, Saint-Constant, Canada) were studied. They were housed in groups of 2 to 4 individuals, provided food and water *ad libitum*, and maintained on a 12 h light:dark cycle. They were purchased when their body weight was between 22 and 24 g, and they were acclimatized one week prior to the start of the experiments. All procedures were approved by the Committee of Animal Care of *Université Laval* following the guidelines from the Canadian Council on Animal Care (2023-1254) and complied with ARRIVE guidelines.

### Surgery

Surgery took place ten days before the first exposure to HDM. The whole protocol is displayed in Figure 1. Mice were first randomly assigned to one of three groups: orchiectomized and supplemented with the vehicle (n=32), sham orchiectomized and supplemented with the vehicle (n=32), and orchiectomized and supplemented with testosterone (n=32). Bilateral orchiectomy was performed as previously described^25^. Briefly, mice were anaesthetized with isoflurane (4% for induction, and 2% thereafter) followed by a subcutaneous injection of buprenorphine (0.05 mg/kg) and Ringer’s lactate (10 mL/kg). The lack of reflex to a toe pinch was used for confirming proper anaesthesia. Local doses of anaesthetics (7 mg/kg of lidocaine and 3.5 mg/kg of bupivacaine) were then injected superficially into the skin of the scrotum and the nape before shaving and cutting the skin open. The testes were then severed. For the sham orchiectomy, testes were externalized and then replaced into the scrotum. A miniature osmotic pump (alzet^®^ osmotic pump, DURECT Corporation, Campbell, CA) was then implanted under the skin of the nape for a continuous delivery of testosterone or its vehicle (40% 2-hydroxypropyl-β-cyclodextrin + 15% ethanol in saline 0.9%). More specifically, the 200 µL pump (model 2004), capable of delivering 0.25 µL/day for 28 days, was used. All mice that had undergone the sham orchiectomy and half of mice that had undergone orchiectomy were implanted with the pump containing the vehicle. The other half of mice that had undergone orchiectomy were implanted with a pump containing testosterone that was delivering 0.25 mg/day/30 g of body weight. After the surgery, mice received a subcutaneous injection of meloxicam (1 mg/kg). They were then inspected closely for the five subsequent days, and the surgical clips on the scrotum and the nape were removed seven days after surgery.

**Figure 1.**
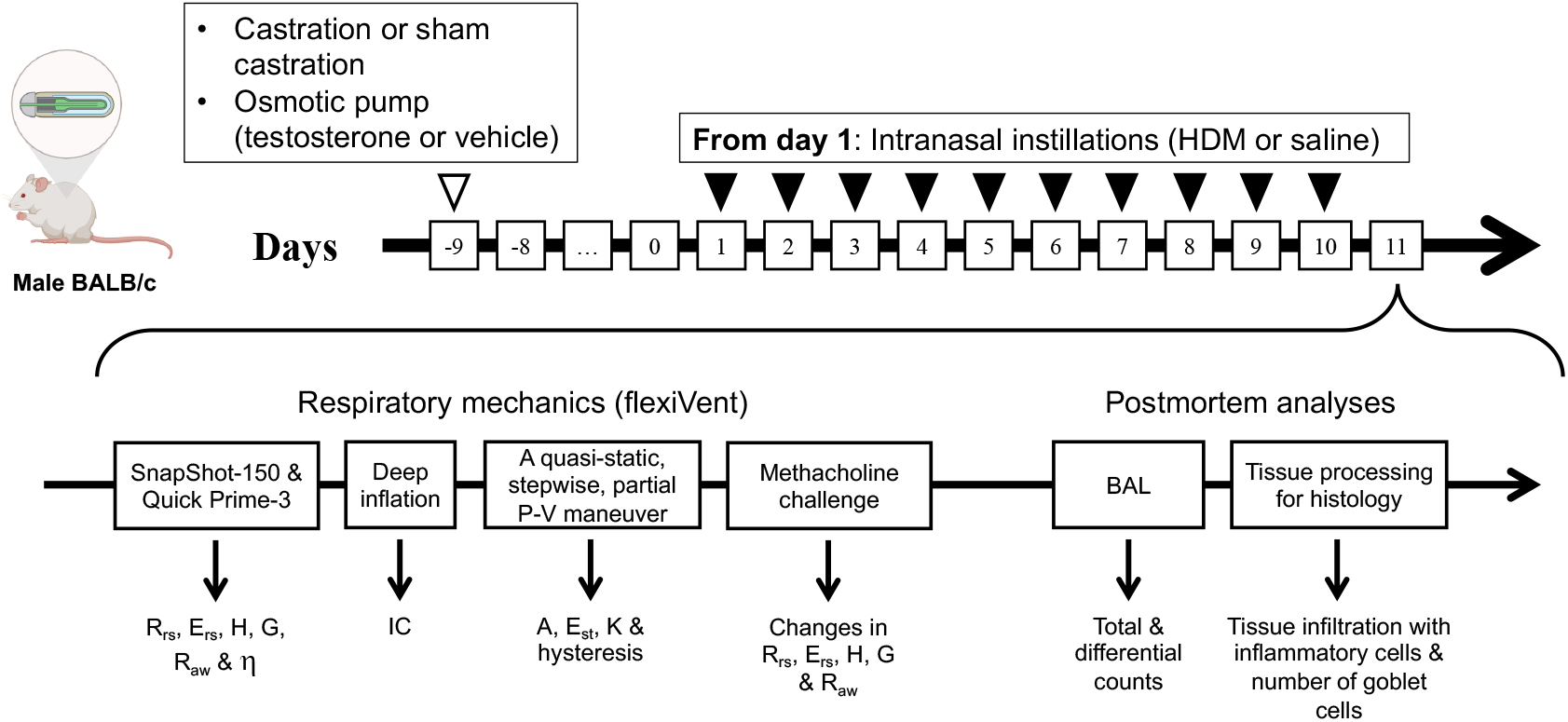
Experimental protocol. See Methods for further details. Abbreviations: A, parameter A of Salazar-Knowles equation; BAL, bronchoalveolar lavages; E_rs_, respiratory system elastance; E_st_, quasi-static elastance; G, tissue resistance; H, tissue elastance; HDM, house dust mite; IC, inspiratory capacity; K, parameter K of Salazar-Knowles equation; P-V, pressure-volume; R_aw_, airway resistance; R_rs_, respiratory system resistance; η (eta), hysteresivity

### Experimental asthma

Experimental asthma was induced by repeated exposures to HDM (Figure 1). The exposure occurred once daily for 10 consecutive days as previously described^23,29,30^. Half of the mice in each group described above (n=16), were exposed to either 25 μL of saline or 25 μL of 2 mg/mL of HDM extract (*D. pteronyssinus, lot number* 428224; Greer, Lenoir, NC) through an intranasal instillation under isoflurane anesthesia. The endotoxin concentration was 19.1 EU per mg of HDM extract. All measurements were made the day after the last exposure.

### Respiratory mechanics

#### Experimental set-up and mechanical ventilation

Mice were first anesthetized and put under general analgesia using ketamine (100 mg/kg) and xylazine (10 mg/kg). Their body weight was measured and they were tracheotomized and connected to the flexiVent (FX Module 2, SCIREQ, Montreal, QC, Canada) through an 18-gauge cannula in a supine position. A surgical thread was tied around the trachea to seal it against the inserted cannula and avoid air leakage. Mice were ventilated mechanically at a tidal volume of 10 mL/kg with an inspiratory-to-expiratory time ratio of 2:3 at a breathing frequency of 150 breaths/min and with a positive end-expiratory pressure (PEEP) of 3 cmH_2_O. Once the ventilation was underway, mice were paralyzed by injecting 100 and 300 µL of pancuronium bromide (0.12 mg/mL) intramuscularly and intraperitoneally, respectively, to avoid spontaneous breathing during the procedure. The maneuvers described next were all measured with the flexiVent in the order illustrated in Figure 1.

#### Deep inflation maneuvers

Mice were subjected to two deep inflation maneuvers (also called recruitment maneuvers). The latter consists of a ramp-style inflation of the lungs from 3 to 40 cmH_2_O over 3 s followed by a 3-s hold at 40 cmH_2_O. The amount of air getting into the lungs during this 6-s maneuver is considered the inspiratory capacity (IC).

#### Oscillometry

Two oscillometric signals were then used to probe respiratory mechanics. One is called the SnapShot-150. It is an input flow signal made of a single sine wave oscillation at 2.5 Hz that allows the calculation of resistance (R_rs_) and elastance (E_rs_) of the respiratory system based on the linear single-compartment model^31^. The other one is called the Quick Prime-3. It is an input flow signal made of 13 sine waves of mutually prime frequencies with different amplitudes and phases, allowing the impedance of the respiratory system to be calculated from the resulting output pressure signal^32^. The impedance was then analyzed using a computational model called the constant phase model to calculate three parameters^33^: 1-Airway resistance (R_aw_), which reflects the resistance to airflow in conducting airways, although it can sometimes be influenced by the chest wall^34-36^; 2-tissue resistance (G), which reflects the tissue resistance of the lungs and the chest wall^34-36^ but is also sensitive to small airway narrowing heterogeneity^37^; and 3-tissue elastance (H), which reflects the elastance of the lungs and is thus sensitive to both the accessible (*i*.*e*., reachable from the mouth) volume of the lungs and the tissue stiffness of the lungs and the chest wall^34,35^. The hysteresivity (η), which is the ratio of G over H, was also determined.

#### Partial pressure-volume (P-V) maneuver

The mice were then subjected to a quasi-static, stepwise, pressure-controlled, partial P-V maneuver (hereinafter called the partial P-V maneuver). It was done as previously described^38,39^. Briefly, it consists of sequentially inflating the lungs through eight steps of increasing pressure (3 to 40 cmH_2_O) and then deflating it through eight steps of decreasing pressure (40 to 3 cmH_2_O). The entire maneuver lasts 16 s. Volume changes at the different holding pressures are recorded and then plotted to form the inflation and deflation limbs of the P-V loop. The descending limb of the P-V loop is then fitted to the Salazar-Knowles equation^40^: V = A -B*e*^-KP^. V and P are the measured volume and pressure, respectively. A, B and K are parameters. The parameter A represents the asymptote on the volume axis. It provides an estimate of the inspiratory capacity. The parameter B (not shown in the present study) represents the difference between A and the extrapolated volume at which pressure would cross zero. Finally, the parameter K is an exponent describing the curvature of the descending limb. It is considered a volume-independent indicator of compliance of the respiratory system^39^. Hysteresis and quasi-static elastance (E_st_) were also extracted from the P-V loop. Hysteresis is the area enclosed by the P-V loop. E_st_ was calculated from the fit of the Salazar-Knowles equation. More precisely, it is the inverse of the fit’s slope at 5 cmH_2_O.

#### The methacholine response

Mice were then subjected to a multiple-concentration challenge with methacholine. The concentrations used were 0, 3, 10, 30, and 100 mg/mL, all diluted in phosphate-buffered saline (PBS). They were nebulized at 5-min intervals. For each concentration, the nebulizer for small particle size (Aeroneb Lab, Aerogen Inc, Galway, Ireland) was operating for a duration of 10 s at a duty cycle of 50% under regular ventilation. Two lung recruitment maneuvers to 30 cmH_2_O were performed before the start of the methacholine challenge and after each subsequent concentration to avoid alveolar collapse, which is in line with optimized protocols of mechanical ventilation^41^.

To monitor the changes in respiratory mechanics during the methacholine challenge, the two oscillometric perturbations described above (*i*.*e*., the SnapShot-150 and the Quick Prime-3) were used. They were each actuated twice at baseline in an alternating fashion. They were then each actuated 12 times, again in an alternating fashion, after each methacholine concentration, starting 27 s after the end of nebulization. Five to seven seconds of tidal breathing was intercalated between each actuation to avoid desaturation. The changes in oscillometric readouts described above, namely R_rs_, E_rs_, H, G, and R_aw_, from their baseline values to their peak values following each concentration of methacholine were used to trace the concentration-response curve. The methacholine response was then quantified by measuring the area under the curve for each oscillometric readout.

### Bronchoalveolar lavages (BAL)

Mice were disconnected from the flexiVent and one mL of PBS was injected into the lungs through the trachea and aspirated to recover the BAL. This was repeated three times and the recovered BAL were pooled. The total volume was recorded and centrifuged at 500 x g during five min. The supernatant was discarded and the pellet was resuspended in 200 μL of PBS for mice exposed to saline and 500 μL for mice exposed to HDM. Total cells in BAL were stained with crystal violet and counted using a hemacytometer. Seventy-five thousand cells were also cytospun and stained with modified May-Grünwald Giemsa to count the number of macrophages, lymphocytes, neutrophils and eosinophils. Their relative proportions were then expressed in percentage.

### Euthanasia and tissue collection

Mice were euthanized by exsanguination after the last flexiVent measurement. The blood was recovered and centrifuged at 2500g at 4°C for 10 min. The supernatant then was collected and stored at -20°C until further processed. The left lung was also excised and immersed in formalin until further processed for histology.

### Histology

Histology was performed in eight mice per group. Histologic alterations seen in this specific model of asthma were previously characterized^23,29^. It was repeated herein in a subgroup of mice to confirm the establishment of experimental asthma. Briefly, after 24 h of fixation in formalin, the lung lobe was progressively dehydrated by upraising the ethanol concentration. It was then embedded in paraffin and cut transversally in 5 μm-thick sections. Sections were deposited on microscopic slides and stained with hematoxylin and eosin (H&E) or periodic acid-Schiff (PAS) with alcian blue. They were then scanned with a NanoZoomer Digital scanner (Hamamatsu photonics, Bridgewater, NJ, USA) at 40X.

H&E stain was performed to evaluate the infiltration of inflammatory cells within the lung tissue. Fifteen non-overlapping photomicrographs (1440 x 904 pixels) from four non-contiguous lung sections were blindly scored from zero (no inflammation) to five (very severe inflammation) by one observer. The scores from each of the 15 photomicrographs were averaged to obtain one value per mouse. The stain with periodic acid-Schiff (PAS) and alcian blue was performed to count goblet cells. All airways cut transversally in three non-contiguous lung sections were analyzed, representing seven to 19 airways per mouse (average of 11.9 ± 3.0). The numbers are expressed in mm of basement membrane length.

### Blood concentration of testosterone

The concentration of testosterone in the serum was quantified by ELISA, as per the manufacturer’s instruction (Mouse/Rat Testosterone ELISA Kit, ab285350, Abcam).

### Statistics

Individual data are presented, together with medians ± interquartile range (IQR). Results were first checked for normality. Since the results of at least one group for each readout were not normally distributed, non-parametric analyses were used throughout. Kruskal-Wallis test was used to compare the serum level of testosterone between groups. If significant, it was followed by Dunn’s multiple comparisons test. Two-factor aligned-rank transform (ART) ANOVAs were then used to assess the effect of testosterone (three levels), experimental asthma (saline *vs*. HDM), and their interaction on readouts of inflammation and respiratory mechanics. When the effect of testosterone was significant, it was followed by contrast tests to determine which level of testosterone differed from the others. All statistical analyses were performed with Prism (version 10.6.0, GraphPad, San Diego, CA), except two-factor ART ANOVAs, which were done in R using the ARTool package (version 0.11.1). Differences with a p<0.05 were considered statistically significant.

## RESULTS

### Testosterone levels

The distinct circulating levels of testosterone were confirmed at day 11, one day after the last HDM exposure. The results are depicted in Figure 2. Orchiectomy and the supplementation with testosterone greatly influenced the level of circulating testosterone (p<0.0001). All pairwise comparisons were also statistically significant (p≤0.0008). Orchiectomized mice implanted with the osmotic pump containing the vehicle were almost devoid of testosterone (0.9 ± 0.6 ng/mL) with a spread ranging from 0.06 to 1.94 ng/mL. Orchiectomized mice implanted with the osmotic pump containing testosterone had the highest levels (73.5 ± 67.1 ng/mL) with a large spread ranging from 6.4 to 264.7 ng/mL. Sham orchiectomized mice implanted with the osmotic pump containing the vehicle had intermediate levels of circulating testosterone (15.2 ± 14.8 ng/mL), also with a large spread ranging from 0.7 to 54.1 ng/mL. The levels of testosterone in these three distinct groups are hereinafter qualified as subphysiologic, physiologic, and supraphysiologic.

**Figure 2.**
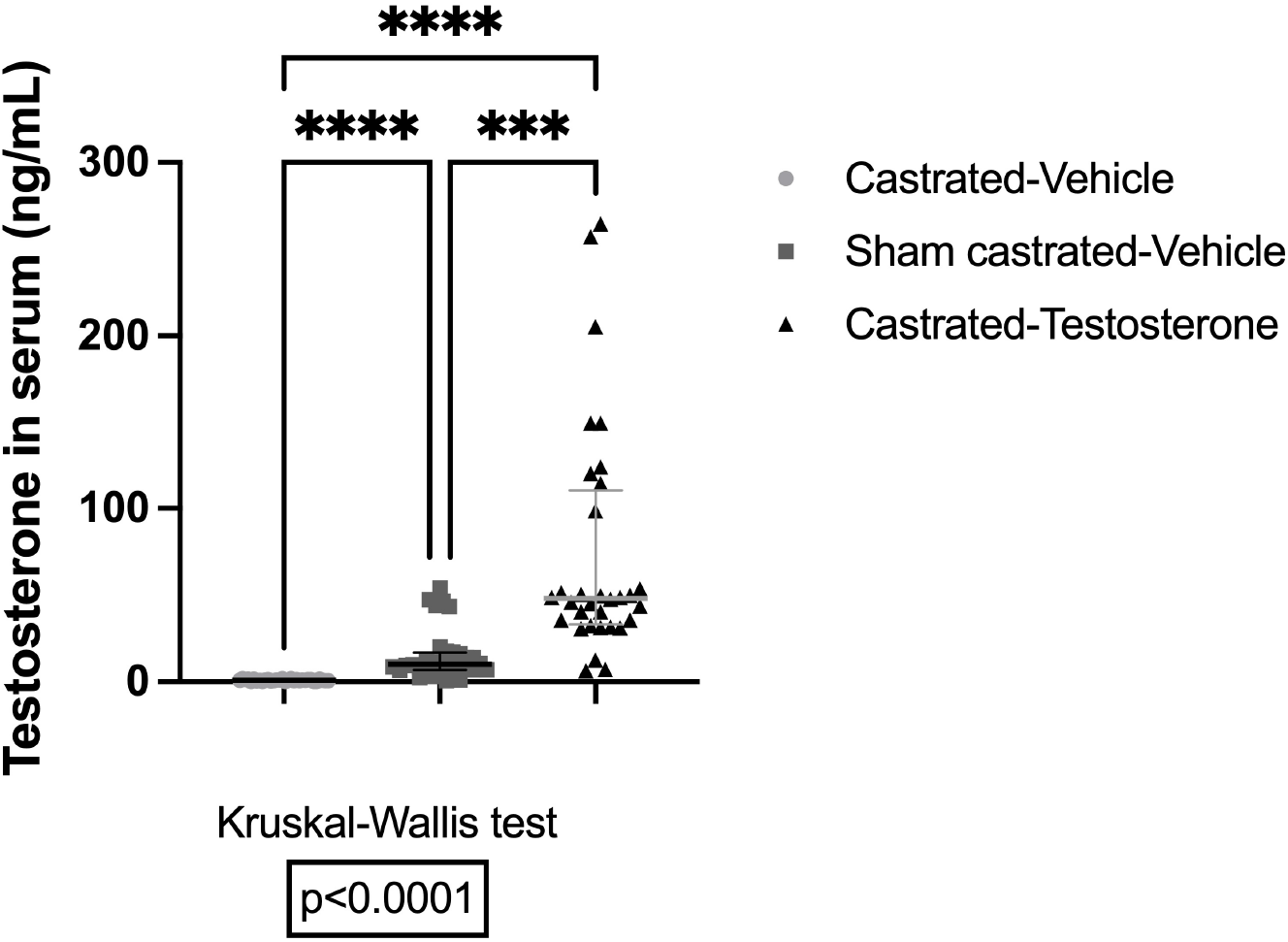
Serum levels of testosterone in castrated mice supplemented with the vehicle (circles), sham castrated mice supplemented with the vehicle (squares), and castrated mice supplemented with testosterone (triangles). The result of the Kruskal-Wallis test is depicted underneath the graph. Dunn’s multiple comparisons test was then used for comparing between different pairs of groups. *** and **** indicate p<0.001 and <0.0001, respectively. N=32

### Body weight

The body weight is depicted in Figure 3. The body weight was greatly influenced by the level of testosterone (p<0.0001) but not by experimental asthma (p=0.62). Mice with subphysiologic levels of testosterone were the lightest (24.8 ± 0.7 g) with a spread ranging from 23.6 to 26.2 g. Mice with supraphysiologic levels of testosterone were the heaviest (27.9 ± 0.9 g) with a spread ranging from 26.0 to 30.9 g. Mice with physiologic levels of testosterone had intermediate body weights (27.2 ± 1.1 g) with a spread ranging from 25.0 to 29.5 g.

**Figure 3.**
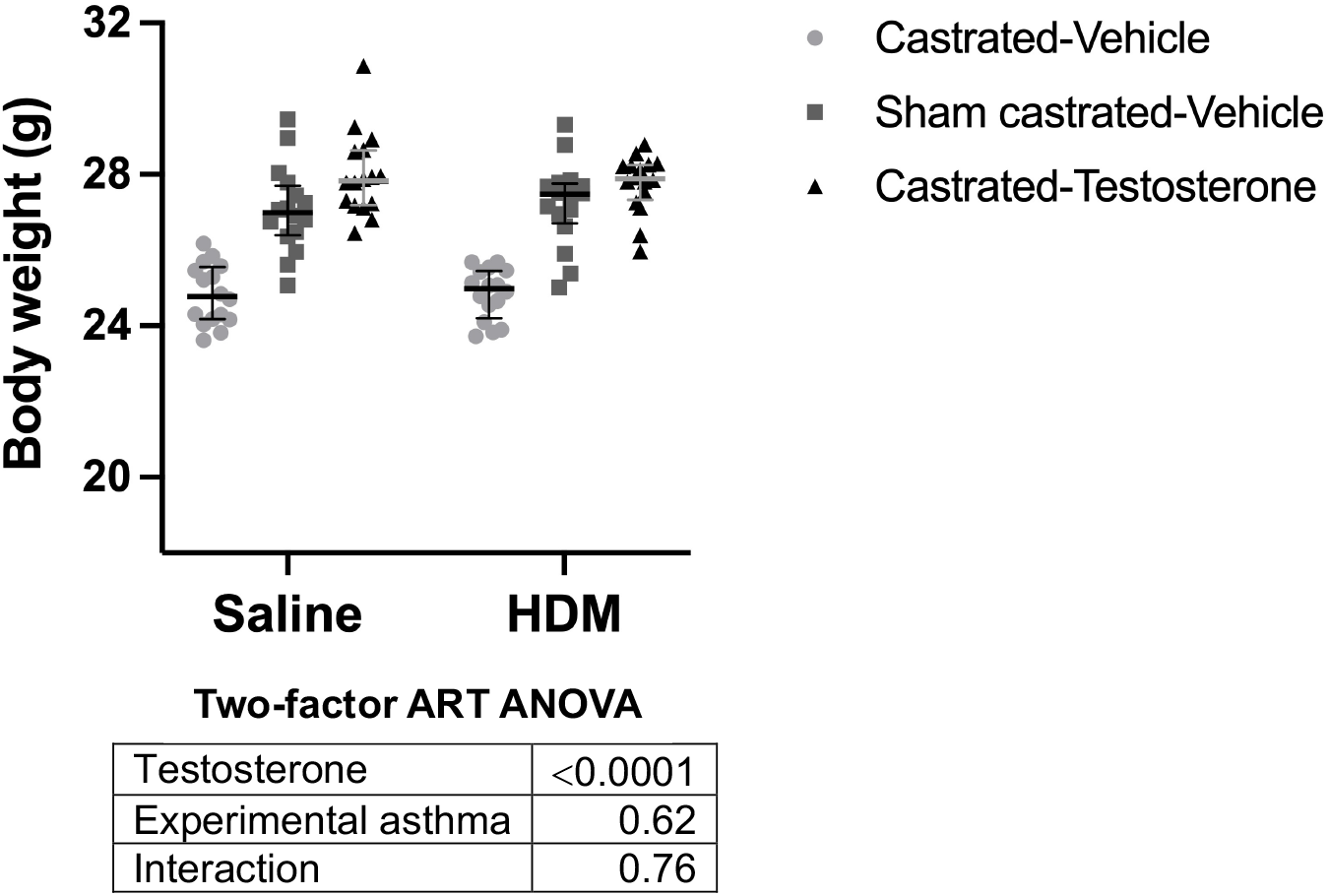
Body weight in castrated mice supplemented with the vehicle (circles), sham castrated mice supplemented with the vehicle (squares), and castrated mice supplemented with testosterone (triangles). The results of the two-factor ART ANOVA are depicted underneath the graph. N=16

### Inflammatory cells in bronchoalveolar lavages (BAL)

Total cell counts in BAL are depicted in Figure 4. Cell counts were influenced by the level of testosterone (p=0.0005) and increased by experimental asthma (p<0.0001). There was also a significant interaction between testosterone levels and experimental asthma (p=0.0003), indicating that testosterone was decreasing cell counts in mice exposed to house dust mite (HDM) but not in mice exposed to saline. Similar statistics were obtained when analyzing cell counts per mL of BAL instead of total cell counts (testosterone level, p=0.001; experimental asthma, p<0.0001; and interaction, p=0.0009).

**Figure 4.**
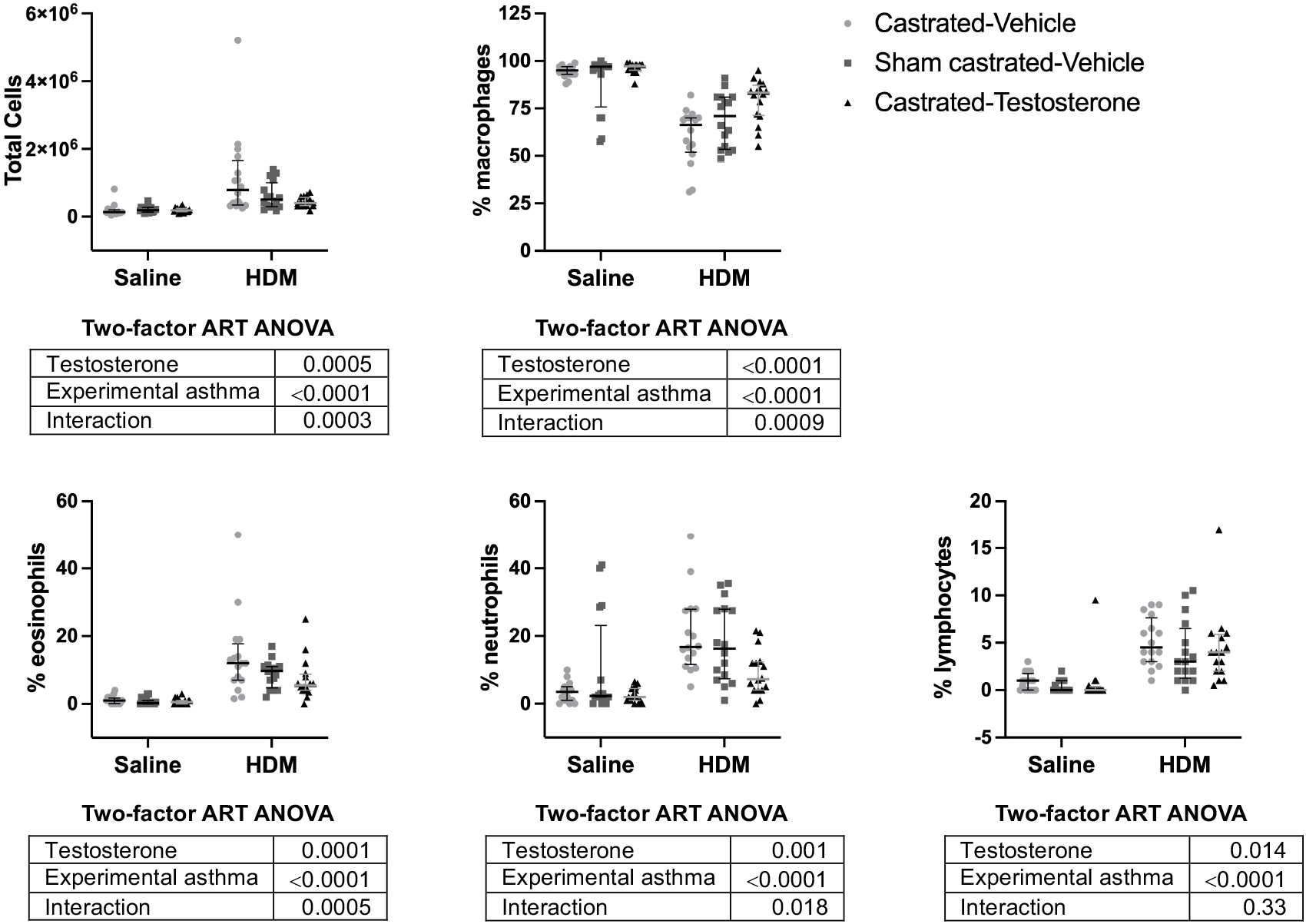
Total cell counts, and the percentage of macrophages, eosinophils, neutrophils and lymphocytes in bronchoalveolar lavages of castrated mice supplemented with the vehicle (circles), sham castrated mice supplemented with the vehicle (squares), and castrated mice supplemented with testosterone (triangles). The results of the two-factor ART ANOVA are depicted underneath each graph. N=16

Differential cell counts in BAL are also depicted in Figure 4. The percentage of macrophages was influenced by the level of testosterone (p<0.0001) and decreased by experimental asthma (p<0.0001). There was also a significant interaction between testosterone levels and experimental asthma (p=0.0009), indicating that testosterone was increasing the percentage of macrophages in mice exposed to HDM but not in mice exposed to saline. The percentage of eosinophils, neutrophils and lymphocytes were influenced by the level of testosterone (p=0.0001, p=0.001 and 0.014, respectively) and increased by experimental asthma (p<0.0001). For eosinophils and neutrophils, there was also a significant interaction between testosterone levels and experimental asthma (p=0.0005, p=0.018, respectively), indicating that testosterone was decreasing the percentage of eosinophils and neutrophils to a greater extent in mice exposed to HDM than in mice exposed to saline. The interaction between testosterone levels and experimental asthma for lymphocytes was not significant (p=0.33).

### Histology

Histology is depicted in Figure 5. The tissue infiltration with inflammatory cells was greatly influenced by the level of testosterone (p=0.002) and increased by experimental asthma (p<0.0001). There was also a significant interaction between testosterone levels and experimental asthma (p=0.040), indicating that testosterone was decreasing inflammatory cell infiltration in mice exposed to HDM to a greater extent than in mice exposed to saline. Similarly, the number of goblet cells was influenced by the level of testosterone (p=0.011) and increased by experimental asthma (p<0.0001). There was no significant interaction between testosterone levels and experimental asthma (p=0.074).

**Figure 5.**
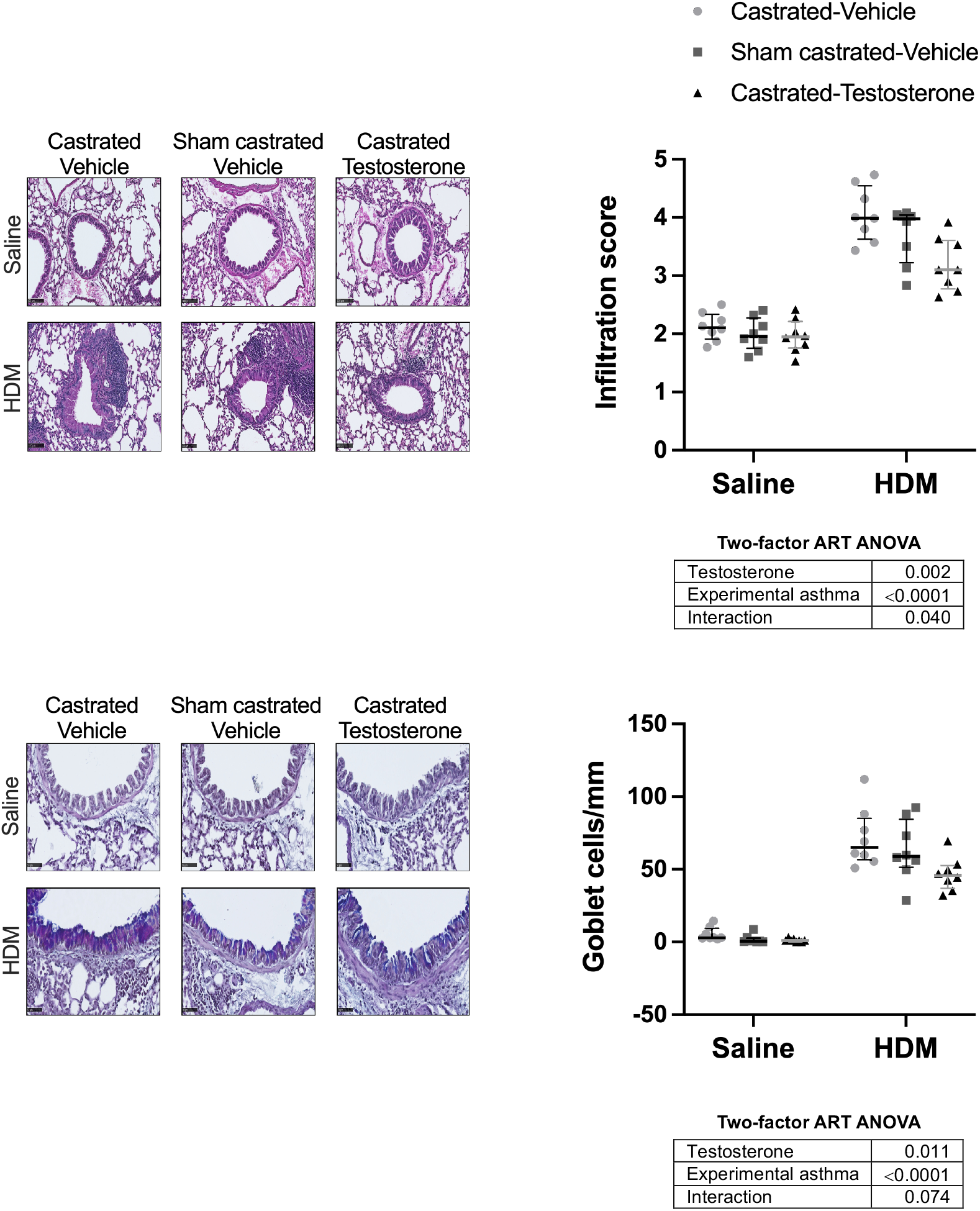
Histologic analyses of tissue infiltration with inflammatory cells (upper panels) and the number of goblets cells per mm of basement membrane (lower panels) in castrated mice supplemented with the vehicle (circles), sham castrated mice supplemented with the vehicle (squares), and castrated mice supplemented with testosterone (triangles). A representative photomicrograph in each group are shown on the left and the quantification in all mice are shown on the right. The results of the two-factor ART ANOVA are depicted underneath each graph. N=16

### Baseline respiratory mechanics

Baseline (*i*.*e*., prior to the methacholine challenge) parameters of respiratory mechanics measured by oscillometry are depicted in Figure 6. Respiratory system resistance (R_rs_), respiratory system elastance (E_rs_), tissue resistance (G), and hysteresivity (η) were significantly affected by testosterone (p=0.009, p=0.001, p<0.0001, and p<0.0001, respectively). In most cases, values were higher in mice with subphysiologic levels of testosterone compared to mice with physiologic levels of testosterone (R_rs_, p=0.007; E_rs_, p=0.058; G, p=0.0003; and η, p<0.0001) and mice with supraphysiologic levels of testosterone (R_rs_, p=0.100; E_rs_, p=0.0006; G, p=0.0004; and η, p=0.002). Tissue elastance (H) and airway resistance (R_aw_) were not affected by testosterone. None of the oscillometric parameters were affected by experimental asthma at baseline.

**Figure 6.**
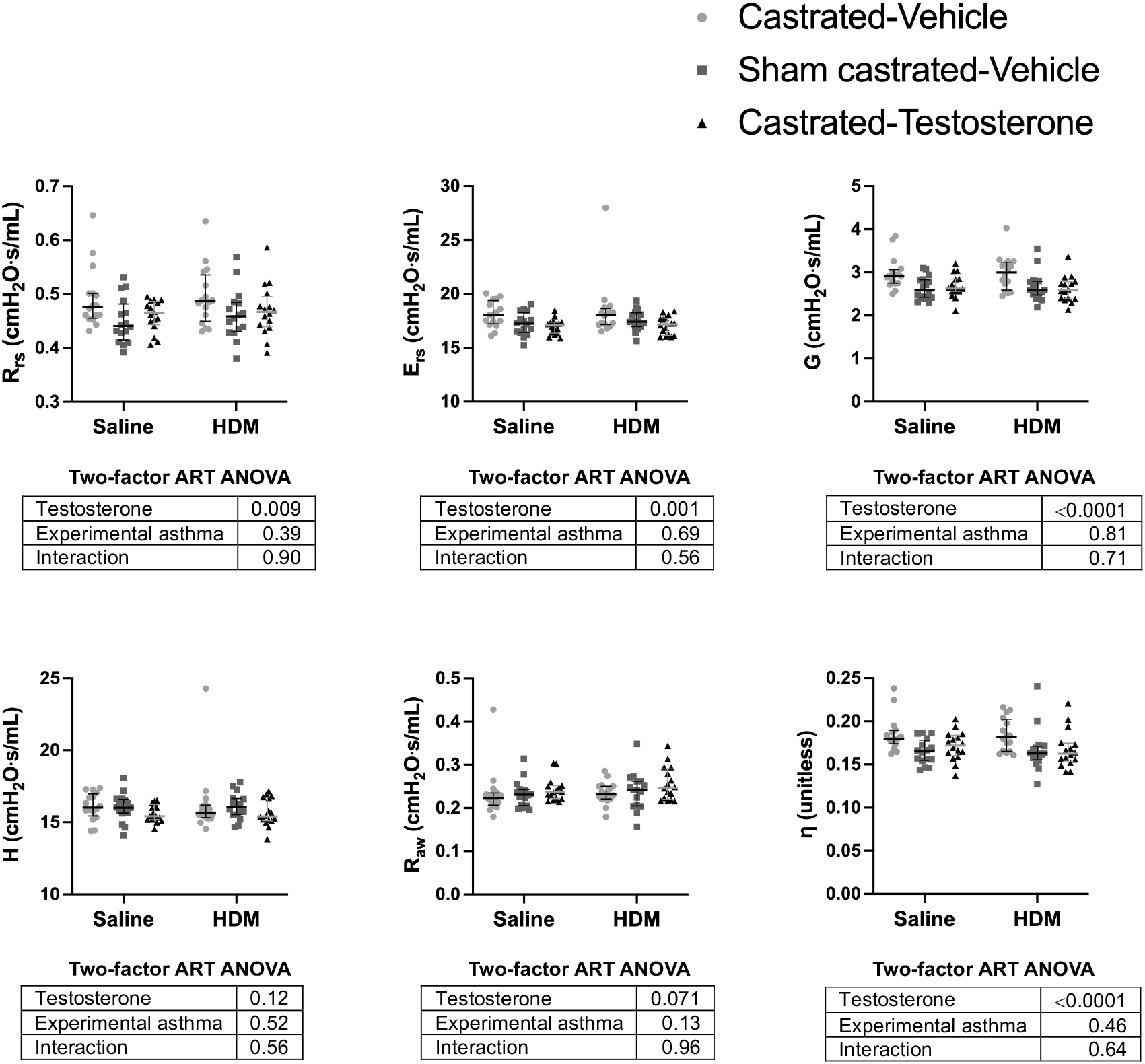
Baseline respiratory mechanics measured by oscillometry. Depicted are system resistance (R_rs_), respiratory system elastance (E_rs_), tissue resistance (G), tissue elastance (H), airway resistance (R_aw_), and hysteresivity (η) for castrated mice supplemented with the vehicle (circles), sham castrated mice supplemented with the vehicle (squares), and castrated mice supplemented with testosterone (triangles). The results of the two-factor ART ANOVA are depicted underneath each graph. N=16

Baseline parameters of respiratory mechanics measured by the partial pressure-volume (P-V) maneuver are depicted in Figure 7. The parameter A of Salazar-Knowles equation (a proxy for the inspiratory capacity), quasi-static elastance (E_st_), and hysteresis (the area enclosed by the P-V loop, which is an indicator of resistive forces) were affected by testosterone (all p<0.0001). Values of the parameter A and hysteresis increased and values of E_st_ decreased with the level of testosterone, all pairwise comparisons between the three levels of testosterone being statistically significant (p≤0.025). The parameter K of Salazar-Knowles equation was not significantly affected by testosterone (p=0.052). Pertaining to experimental asthma, E_st_ increased and K decreased with HDM exposure (p=0.005 and 0.0008, respectively). The parameter A and hysteresis were not affected by experimental asthma. The results of inspiratory capacity (IC) were virtually identical to the ones of the parameter A (data not shown).

**Figure 7.**
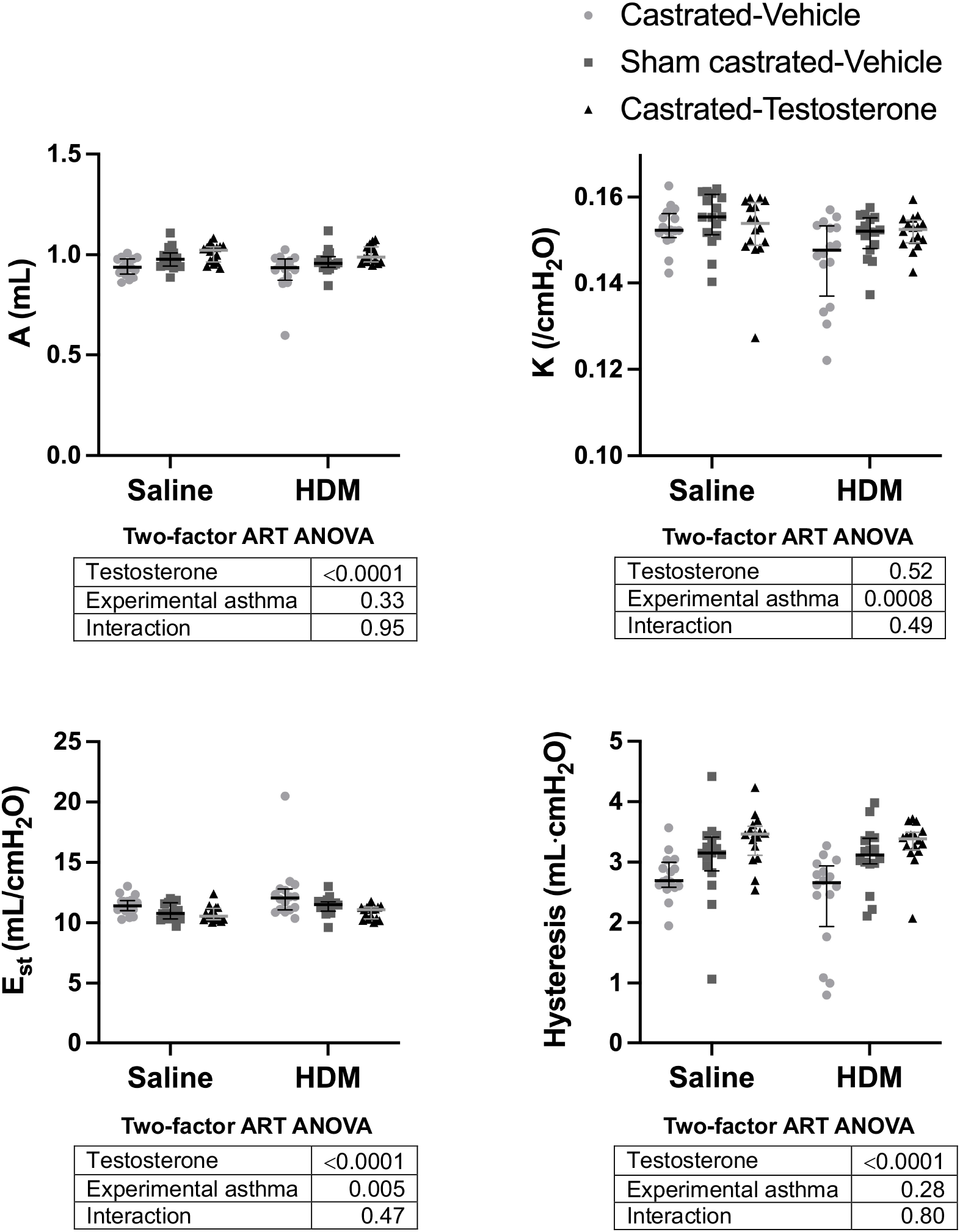
Baseline respiratory mechanics measured from the partial P-V maneuver. Depicted are two parameters from the Salazar-Knowles equation (A & K), quasi-static elastance (E_st_), and hysteresis for castrated mice supplemented with the vehicle (circles), sham castrated mice supplemented with the vehicle (squares), and castrated mice supplemented with testosterone (triangles). The results of the two-factor ART ANOVA are depicted underneath each graph. N=16

### The methacholine response

The response to nebulized methacholine is depicted in Figure 8. Testosterone significantly increased the methacholine response. This was true when the methacholine response was monitored by measuring the changes in R_rs_ (p=0.037), E_rs_ (p=0.032), G (p=0.048), and H (p=0.039), but not R_aw_ (p=0.108). Experimental asthma also significantly increased the response to methacholine regardless of which parameter was chosen to monitor the response (R_rs_, p<0.0001; E_rs_, p<0.0001; G, p<0.0001; H, p<0.0001; and R_aw_, p=0.002), confirming hyperresponsiveness.

**Figure 8.**
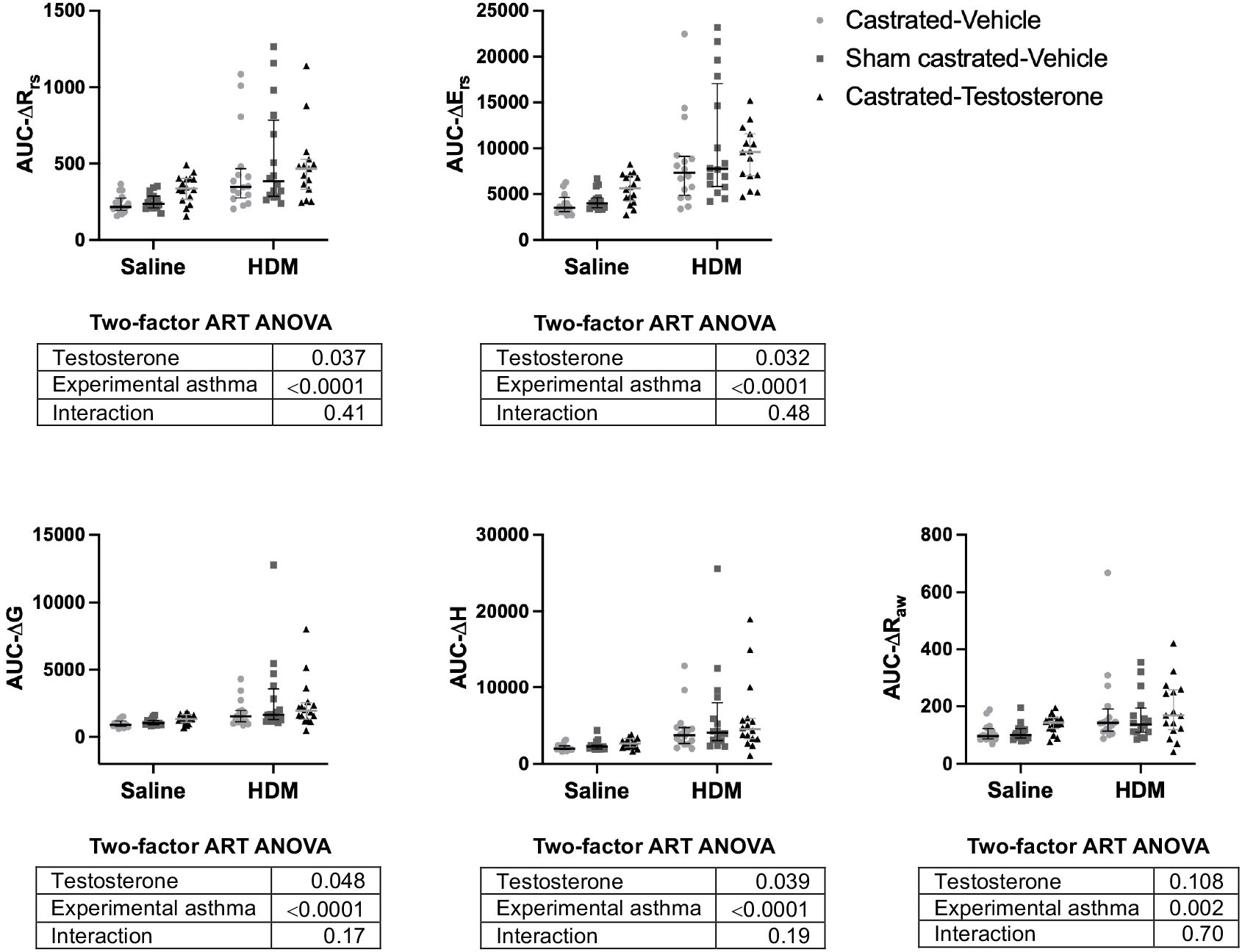
The methacholine response measured from the area under the curve of the concentration-response curves for the changes in respiratory system resistance (AUC-ΔR_rs_), respiratory system elastance (AUC-ΔE_rs_), tissue resistance (AUC-ΔG), tissue elastance (AUC-ΔH), and airway resistance (AUC-ΔR_aw_) in castrated mice supplemented with the vehicle (circles), sham castrated mice supplemented with the vehicle (squares), and castrated mice supplemented with testosterone (triangles). The results of the two-factor ART ANOVA are depicted underneath each graph. N=16

## DISCUSSION

The combination of orchiectomy and the implantation of osmotic pumps was successful for generating three groups of mice with distinct levels of circulating testosterone. The sustained delivery of testosterone through the osmotic pump provided a clear edge compared to previous studies using repeated boluses that presumably yielded transient peaks in circulating concentrations^9,13,16^. As such, our results are not expected to perfectly align with others. Our results demonstrate that the mouse body weight was commensurate to testosterone levels, suggesting that testosterone promotes weight gain (presumptively through growth and anabolism). Baseline respiratory mechanics were also significantly affected by testosterone. In most cases, mice with a subphysiologic level of testosterone were worse compared to the other two groups. Most facets of inflammation were also affected by testosterone. In all cases, testosterone exerted an anti-inflammatory effect, decreasing total cell counts in BAL, the % of eosinophils and neutrophils in BAL, the tissue infiltration with inflammatory cells, and the number of goblet cells in the epithelium, and increasing the % of macrophages in BAL. This anti-inflammatory effect was mainly seen in inflamed mice exposed to HDM, as mice exposed to saline had no inflammation. Finally, testosterone significantly increased the response to nebulized methacholine.

Our results are consistent with a compendium of recent publications suggesting that testosterone decreases inflammation in murine models of asthma^7-17^. Inasmuch as mice and rat relate to humans, this anti-inflammatory effect alone may explain the protective role of testosterone in asthma. Yet, how testosterone exerts its anti-inflammatory effects is still not clear. In the present study, all measured aspects of inflammation were decreased (Figures 4 & 5). Thus, it seems that testosterone exerts a broad role, buffering the overall inflammation. It is possible that testosterone acts upstream on a specific orchestrating mechanism that controls inflammation in general. Alternatively, testosterone may act downstream on many features of inflammation. Previous studies have pointed towards an inhibition of allergic sensitization/inflammation and airway remodeling^7,9,17^ by; 1-suppressing IL-4 and IL-17A production^7^; 2-negatively controlling group 2 innate lymphoid cells (ILC2s)^8,11^; 3-promoting the suppressive function of regulatory T cells (Tregs)^14^; 4-acting directly on CD4^+^ T cells and downregulating cytokine production^17^; 5-facilitating T-helper (Th) lymphocytes differentiation into Th1^13^; 6-promoting the inhibition of Th2 cell differentiation^16^; and 7-restricting glutaminolysis and suppressing Th17 cells^15^. Some of its cell-specific actions may also promote experimental asthma. For example, despite decreasing the overall allergic lung inflammation, testosterone enhanced the polarization of alveolar macrophages into M2, which was required in males for a full-blown allergic lung inflammation^10^. More studies of this kind, scrutinizing the many important components of inflammation, are warranted to unveil the intricacies by which testosterone negatively and positively influences the establishment of inflammation in experimental asthma.

Our study was more specifically designed to provide a fine-grained analysis of testosterone on respiratory mechanics, and to shed light on seemingly inconsistent observations pertaining to the methacholine response. The first noticeable findings were seen in mice with a subphysiologic level of testosterone, showing aberrant respiratory mechanics at baseline. More precisely, altered mechanics in this group were shown by increases in R_rs_, E_rs_, G, η, and E_st_, as well as decreases in A, IC, and hysteresis.

The mouse size (*i*.*e*., body weight) may be contributing to these differences, as lung mechanics is inversely proportional to size, with lower resistance and elastance in larger subjects. However, we are not sure whether the approximately 3-g difference between the lightest and heaviest groups (Figure 3) would entirely explain the differences in mechanics. One argument against the body size involvement is the significant difference in hysteresivity (Figure 6). Hysteresivity is an intensive property, meaning that it should not be affected by the amount of tissue being probed but only by changes in the very nature of the tissue^42,43^. Hence, a decreased lung size (as expected from a smaller mouse) should increase both G and H in the same proportion, leaving hysteresivity (*i*.*e*., the ratio of G on H) unchanged. Since hysteresivity was elevated in mice with a subphysiologic level of testosterone, it implies that the nature of their tissue was different. The lack of differences in H (Figure 6), a readout that is sensitive to lung size (and therefore body size), is also another argument against the body size involvement in baseline differences in mechanics.

The lack of difference in H is also perplexing because E_rs_, which normally matches the results of H, was higher in mice with a subphysiologic level of testosterone compared to the other two groups (Figure 6). The only logical explanation is that we are dealing with a frequency-dependent phenomenon, as H is deduced by probing the lungs across a range of frequencies (1 to 20.5 Hz), whereas E_rs_ is deduced by probing the lungs at a single low frequency (2.5 Hz). Since low frequency oscillations are reputed to travel deeper into the lungs, would that mean that only the most peripheral parts of the lungs were affected in mice with a subphysiologic level of testosterone?

The lack of difference in H also suggests that the increased hysteresivity in mice with a subphysiologic level of testosterone was entirely driven by an elevated G (Figure 6). Interestingly, it is well known that G but not H is affected by small airway narrowing heterogeneity^37^, and that narrowing heterogeneity is a phenomenon that changes mechanics in a frequency-dependent fashion^44^. Heterogeneity also precedes closure^37^, and closure is a dynamic phenomenon typically proceeding from the peripheral to the interior parts of the lungs. Therefore, at the time respiratory mechanics was measured, which was after two recruitment maneuvers, it is possible that narrowing heterogeneity was already perceptible (increasing G) but closure (and the attendant increase in elastance) was only perceived at low frequencies deep into the lungs (increasing E_rs_ but not H). The small amount of peripheral closure (with its attendant air trapping) would also explain why less air got into the lungs during large-amplitude maneuvers (*i*.*e*., deep inflation maneuvers and the partial P-V maneuver) and, thereby, explain the decreases in IC and A in mice with a subphysiologic level of testosterone (Figure 7). Taken together, these results suggest that the near lack of testosterone in mice worsens respiratory mechanics not by affecting body size or the nature of the lung tissue, but by promoting airway narrowing heterogeneity and closure.

The second noticeable finding was the potentiating effect of testosterone on the response to nebulized methacholine (Figure 8). This finding is consistent with previous studies on healthy mice^18,19,25^. The present study thus extends these findings by showing that this effect holds in a dataset involving mice with and without experimental asthma. The mechanisms by which testosterone increases the methacholine response is not fully delineated. One study suggested an indirect mechanism involving the vagal nerves. More precisely, the authors demonstrated that bilateral vagotomy (*i*.*e*., severing vagal nerves running on either side of the trachea) prevents the amplifying effect of testosterone on the methacholine response^19^.

Owing to the great influence of testosterone on inflammation, we were surprised that there was no significant interaction between testosterone and experimental asthma on the methacholine response. Inflammation is expected to increase the methacholine response^45^. In fact, inflammation is considered liable for the emergence of hyperresponsiveness, in great part by increasing the propensity for small airway narrowing heterogeneity and closure^37,46-54^. The buffering effect of testosterone on inflammation was thus expected to reduce the response to methacholine in mice with experimental asthma. Previous investigations have actually suggested an attenuating effect of testosterone on the methacholine response in mice with experimental asthma^7,9,15^. The lack of interaction in our dataset rather suggests that the potentiating effect of testosterone on the methacholine response was seen in mice with and without experimental asthma. It probably comes down to some sort of balance between the level of airway smooth muscle (ASM) contraction and the level of inflammation. Since there was no inflammation in saline-exposed mice, testosterone probably enhanced the methacholine response in this group by increasing the contraction of the ASM. If this effect also occurred in mice with experimental asthma, it should have acted synergistically with inflammation to further amplify the methacholine response^45^. The fact that the potentiating effect of testosterone on the methacholine response was similar in mice with experimental asthma suggests that its buffering effect on inflammation must have attenuated the methacholine response. Hence, the HDM-exposed mice with the least inflammation (mice with a supraphysiologic level of testosterone) were not the least responsive to methacholine because the ASM must have been stronger than normal, and the HDM-exposed mice with the worse inflammation (mice with a subphysiologic level of testosterone) were not the most responsive to methacholine because the ASM must have been weaker than normal. Overall, our results do not seem to perfectly align with others, but they are not completely inconsistent neither. With the right doses of testosterone, methacholine, and HDM (perhaps causing a higher level of inflammation with a greater propensity for heterogeneity and closure), one can imagine a scenario where testosterone would mitigate the methacholine response in mice with experimental asthma; *i*.*e*., when the attenuating effect of decreased inflammation on the methacholine response outweighs the potentiating effect of increased ASM contraction. This was presumably the context in which these previous investigators were working^7,9,15^.

Hysteresis, which was long suggested as an indicator of ASM activation^55^, was also elevated by testosterone in the present study (Figure 7). This represents further indirect evidence that testosterone increases the activation of the ASM. Based on the entrenched conviction that ASM contraction contributes to asthma symptoms, increasing the level of ASM activation may sound like something detrimental. However, olden^56,57^ and recent^21,23,58^ evidence rather suggest that a low level of ASM activation in the peripheral parts of the lungs is important to prevent small airway narrowing heterogeneity and closure. This would be supported by the present study, based on the altered respiratory mechanics at baseline seen in mice with a subphysiologic level of testosterone. The presumptive increased activation of the ASM caused by testosterone may also increase the contractile capacity of the ASM through a process called force adaptation^59^, which may then contribute to the increased response to methacholine caused by testosterone. Interestingly, and despite increasing the ASM’s contractile capacity, the process of force adaption was shown to be rather protective against hyperresponsiveness in experimental asthma^58^. The proposed underlying mechanism involves the stiffening of the lung tissue, which then forces the lungs to work more homogeneously and thereby prevents small airway narrowing heterogeneity and closure^58^, which are major causes of hyperresponsiveness in asthma and experimental asthma^37,46-54^. Perhaps a similar mechanism was operating herein, offering an alternative interpretation for the lack of synergistic effect between testosterone and experimental asthma on the methacholine response.

## Conclusion

Testosterone decreases allergic lung inflammation in male BALB/c mice with experimental asthma. This anti-inflammatory effect alone may explain its seemingly beneficial effect in asthma. Yet, several evidence presented herein, as well as from previous studies^18,19,25^, indicate that testosterone increases the level of ASM activation. In a more severe inflammatory context, it is possible that this increased contractility of the ASM stiffens the lung tissue and thereby mitigates hyperresponsiveness by preventing narrowing heterogeneity and closure^58^, which may turn out to be also protective in asthma. We therefore argue that the salutary effect of testosterone in asthma stems from a two-birds-with-one-stone phenomenon, decreasing inflammation and increasing ASM activation. More studies will obviously be required to ascertain the latter.

## AUTHOR’S CONTRIBUTIONS

All authors edited the manuscript, and read and approved the final manuscript.

CH, MB, DM, VJ and YB contributed to the development of the experimental design.

CH, MB, ARR, LC, LG, and MJB performed laboratory experiments.

CH, MB, MJB, and YB analyzed the data.

YB wrote the first version of the manuscript.

## DATA AVAILABILITY STATEMENT

All data are shown and can be made available in other formats from the corresponding author on reasonable request.

## REFERENCES

1 Fuseini, H. & Newcomb, D. C. Mechanisms Driving Gender Differences in Asthma. Curr Allergy Asthma Rep 17, 19, doi:10.1007/s11882-017-0686-1 (2017).

2 Zein, J. G. & Erzurum, S. C. Asthma is Different in Women. Curr Allergy Asthma Rep 15, 28, doi:10.1007/s11882-015-0528-y (2015).

3 Melgert, B. N., Ray, A., Hylkema, M. N., Timens, W. & Postma, D. S. Are there reasons why adult asthma is more common in females? Curr Allergy Asthma Rep 7, 143–150, doi:10.1007/s11882-007-0012-4 (2007).

4 Kelsey, T. W. et al. A validated age-related normative model for male total testosterone shows increasing variance but no decline after age 40 years. PLoS One 9, e109346, doi:10.1371/journal.pone.0109346 (2014).

5 Bulkhi, A. A., Shepard, K. V., 2nd, Casale, T. B. & Cardet, J. C. Elevated Testosterone Is Associated with Decreased Likelihood of Current Asthma Regardless of Sex. J Allergy Clin Immunol Pract 8, 3029–3035 e3024, doi:10.1016/j.jaip.2020.05.022 (2020).

6 DeBoer, M. D. et al. Effects of endogenous sex hormones on lung function and symptom control in adolescents with asthma. BMC Pulm Med 18, 58, doi:10.1186/s12890-018-0612-x (2018).

7 Fuseini, H. et al. Testosterone Decreases House Dust Mite-Induced Type 2 and IL-17A-Mediated Airway Inflammation. J Immunol 201, 1843–1854, doi:10.4049/jimmunol.1800293 (2018).

8 Laffont, S. et al. Androgen signaling negatively controls group 2 innate lymphoid cells. J Exp Med 214, 1581–1592, doi:10.1084/jem.20161807 (2017).

9 Kalidhindi, R. S. R. et al. Androgen receptor activation alleviates airway hyperresponsiveness, inflammation, and remodeling in a murine model of asthma. Am J Physiol Lung Cell Mol Physiol 320, L803–L818, doi:10.1152/ajplung.00441.2020 (2021).

10 Becerra-Diaz, M., Strickland, A. B., Keselman, A. & Heller, N. M. Androgen and Androgen Receptor as Enhancers of M2 Macrophage Polarization in Allergic Lung Inflammation. J Immunol 201, 2923–2933, doi:10.4049/jimmunol.1800352 (2018).

11 Cephus, J. Y. et al. Testosterone Attenuates Group 2 Innate Lymphoid Cell-Mediated Airway Inflammation. Cell Rep 21, 2487–2499, doi:10.1016/j.celrep.2017.10.110 (2017).

12 Montano, L. M., Espinoza, J., Flores-Soto, E., Chavez, J. & Perusquia, M. Androgens are bronchoactive drugs that act by relaxing airway smooth muscle and preventing bronchospasm. J Endocrinol 222, 1–13, doi:10.1530/JOE-14-0074 (2014).

13 He, Y. et al. Combination of androgen and estrogen improves asthma by mediating Runx3 expression. Int J Med Sci 21, 1003–1015, doi:10.7150/ijms.91253 (2024).

14 Gandhi, V. D. et al. Androgen receptor signaling promotes Treg suppressive function during allergic airway inflammation. J Clin Invest 132, doi:10.1172/JCI153397 (2022).

15 Chowdhury, N. U. et al. Androgen signaling restricts glutaminolysis to drive sex-specific Th17 metabolism in allergic airway inflammation. J Clin Invest 134, doi:10.1172/JCI177242 (2024).

16 Chen, Z. et al. Androgens have therapeutic potential in T2 asthma by mediating METTL3 in bronchial epithelial cells. Int Immunopharmacol 143, 113322, doi:10.1016/j.intimp.2024.113322 (2024).

17 Ejima, A. et al. Androgens Alleviate Allergic Airway Inflammation by Suppressing Cytokine Production in Th2 Cells. J Immunol 209, 1083–1094, doi:10.4049/jimmunol.2200294 (2022).

18 Card, J. W. et al. Gender differences in murine airway responsiveness and lipopolysaccharide-induced inflammation. J Immunol 177, 621–630, doi:10.4049/jimmunol.177.1.621 (2006).

19 Card, J. W. et al. Male sex hormones promote vagally mediated reflex airway responsiveness to cholinergic stimulation. Am J Physiol Lung Cell Mol Physiol 292, L908–914, doi:10.1152/ajplung.00407.2006 (2007).

20 Ekpruke, C. D. et al. Sex-specific alterations in the gut and lung microbiome of allergen-induced mice. Front Allergy 5, 1451846, doi:10.3389/falgy.2024.1451846 (2024).

21 Boucher, M. et al. In mice of both sexes, repeated contractions of smooth muscle in vivo greatly enhance the response of peripheral airways to methacholine. Respir Physiol Neurobiol 304, 103938, doi:10.1016/j.resp.2022.103938 (2022).

22 Boucher, M., Henry, C. & Bossé, Y. Force adaptation through the intravenous route in naive mice. Exp Lung Res 49, 131–141, doi:10.1080/01902148.2023.2237127 (2023).

23 Gill, R., Rojas-Ruiz, A., Boucher, M., Henry, C. & Bossé, Y. More airway smooth muscle in males versus females in a mouse model of asthma: A blessing in disguise? Exp Physiol 108, 1080–1091, doi:10.1113/EP091236 (2023).

24 D’Abreo J J. et al. The effects of e-cigarette use on asthma severity in adult BALB/c mice. Exp Physiol, doi:10.1113/EP092959 (2025).

25 Ganouna-Cohen, G., Khadangi, F., Marcouiller, F., Bossé, Y. & Joseph, V. Additive effects of orchiectomy and intermittent hypoxia on lung mechanics and inflammation in C57BL/6J male mice. Exp Physiol 107, 68–81, doi:10.1113/EP090050 (2022).

26 O’Byrne, P. M. Introduction: Airway hyperresponsiveness in asthma: its measurement and clinical significance. Chest 138, 1S–3S, doi:10.1378/chest.10-0091 (2010).

27 Chapman, D. G. & Irvin, C. G. Mechanisms of Airway Hyperresponsiveness in Asthma: The Past, Present and Yet to Come. Clin Exp Allergy, doi:10.1111/cea.12506 (2015).

28 Wulfsohn, N. L., Politzer, W. M. & Henrico, J. S. Testosterone Therapy in Bronchial Asthma. South African medical journal = Suid-Afrikaanse tydskrif vir geneeskunde 38, 170–172 (1964).

29 Boucher, M., Henry, C., Dufour-Mailhot, A., Khadangi, F. & Bossé, Y. Smooth Muscle Hypocontractility and Airway Normoresponsiveness in a Mouse Model of Pulmonary Allergic Inflammation. Front Physiol 12, 698019, doi:10.3389/fphys.2021.698019 (2021).

30 Boucher, M., Henry, C., Khadangi, F., Dufour-Mailhot, A. & Bossé, Y. Double-chamber plethysmography versus oscillometry to detect baseline airflow obstruction in a model of asthma in two mouse strains. Exp Lung Res 47, 390–401, doi:10.1080/01902148.2021.1979693 (2021).

31 Bates, J. H. Lung mechanics: an inverse modeling approach. (Cambridge University Press, 2009).

32 Bates, J. H., Irvin, C. G., Farré, R. & Hantos, Z. Oscillation mechanics of the respiratory system. Comprehensive Physiology 1, 1233–1272 (2011).

33 Hantos, Z., Daroczy, B., Suki, B., Nagy, S. & Fredberg, J. Input impedance and peripheral inhomogeneity of dog lungs. Journal of applied physiology 72, 168–178 (1992).

34 Sudy, R. et al. Different contributions from lungs and chest wall to respiratory mechanics in mice, rats, and rabbits. J Appl Physiol (1985) 127, 198–204, doi:10.1152/japplphysiol.00048.2019 (2019).

35 Hirai, T., McKeown, K. A., Gomes, R. F. & Bates, J. H. Effects of lung volume on lung and chest wall mechanics in rats. J Appl Physiol (1985) 86, 16–21, doi:10.1152/jappl.1999.86.1.16 (1999).

36 Ito, S., Lutchen, K. R. & Suki, B. Effects of heterogeneities on the partitioning of airway and tissue properties in normal mice. J Appl Physiol (1985) 102, 859–869, doi:10.1152/japplphysiol.00884.2006 (2007).

37 Lutchen, K. R., Hantos, Z., Petak, F., Adamicza, A. & Suki, B. Airway inhomogeneities contribute to apparent lung tissue mechanics during constriction. J Appl Physiol (1985) 80, 1841–1849, doi:10.1152/jappl.1996.80.5.1841 (1996).

38 Robichaud, A. et al. Automated full-range pressure-volume curves in mice and rats. J Appl Physiol (1985) 123, 746–756, doi:10.1152/japplphysiol.00856.2016 (2017).

39 Boucher, M. et al. Effects of airway smooth muscle contraction and inflammation on lung tissue compliance. Am J Physiol Lung Cell Mol Physiol 322, L294–L304, doi:10.1152/ajplung.00384.2021 (2022).

40 Salazar, E. & Knowles, J. H. An Analysis of Pressure-Volume Characteristics of the Lungs. J Appl Physiol 19, 97–104, doi:10.1152/jappl.1964.19.1.97 (1964).

41 Reiss, L. K., Kowallik, A. & Uhlig, S. Recurrent recruitment manoeuvres improve lung mechanics and minimize lung injury during mechanical ventilation of healthy mice. PLoS One 6, e24527, doi:10.1371/journal.pone.0024527 (2011).

42 Bates, J. H., Rincon, M. & Irvin, C. G. Animal models of asthma. Am J Physiol Lung Cell Mol Physiol 297, L401–410, doi:10.1152/ajplung.00027.2009 (2009).

43 Fredberg, J. J. & Stamenovic, D. On the imperfect elasticity of lung tissue. J Appl Physiol (1985) 67, 2408–2419, doi:10.1152/jappl.1989.67.6.2408 (1989).

44 Bates, J. H., Irvin, C. G., Farre, R. & Hantos, Z. Oscillation mechanics of the respiratory system. Compr Physiol 1, 1233–1272, doi:10.1002/cphy.c100058 (2011).

45 Bossé, Y., Riesenfeld, E. P., Pare, P. D. & Irvin, C. G. It’s not all smooth muscle: non-smooth-muscle elements in control of resistance to airflow. Annu Rev Physiol 72, 437–462, doi:10.1146/annurev-physiol-021909-135851 (2010).

46 Dame Carroll, J. R., Magnussen, J. S., Berend, N., Salome, C. M. & King, G. G. Greater parallel heterogeneity of airway narrowing and airway closure in asthma measured by high-resolution CT. Thorax 70, 1163–1170, doi:10.1136/thoraxjnl-2014-206387 (2015).

47 Downie, S. R. et al. Ventilation heterogeneity is a major determinant of airway hyperresponsiveness in asthma, independent of airway inflammation. Thorax 62, 684–689, doi:thx.2006.069682 [pii] 10.1136/thx.2006.069682 (2007).

48 Farrow, C. E. et al. Peripheral ventilation heterogeneity determines the extent of bronchoconstriction in asthma. J Appl Physiol (1985) 123, 1188–1194, doi:10.1152/japplphysiol.00640.2016 (2017).

49 Hardaker, K. M. et al. Predictors of airway hyperresponsiveness differ between old and young patients with asthma. Chest 139, 1395–1401, doi:10.1378/chest.10-1839 (2011).

50 King, G. G. et al. Heterogeneity of narrowing in normal and asthmatic airways measured by HRCT. Eur Respir J 24, 211–218 (2004).

51 Petak, F., Hantos, Z., Adamicza, A., Asztalos, T. & Sly, P. D. Methacholine-induced bronchoconstriction in rats: effects of intravenous vs. aerosol delivery. J Appl Physiol (1985) 82, 1479–1487, doi:10.1152/jappl.1997.82.5.1479 (1997).

52 Chapman, D. G., Berend, N., King, G. G. & Salome, C. M. Increased airway closure is a determinant of airway hyperresponsiveness. Eur Respir J 32, 1563–1569, doi:10.1183/09031936.00114007 (2008).

53 Farrow, C. E. et al. Airway closure on imaging relates to airway hyperresponsiveness and peripheral airway disease in asthma. J Appl Physiol 113, 958–966, doi:10.1152/japplphysiol.01618.2011 (2012).

54 Wagers, S., Lundblad, L. K., Ekman, M., Irvin, C. G. & Bates, J. H. The allergic mouse model of asthma: normal smooth muscle in an abnormal lung? J Appl Physiol 96, 2019–2027 (2004).

55 Vincent, N. J., Knudson, R., Leith, D. E., Macklem, P. T. & Mead, J. Factors influencing pulmonary resistance. J Appl Physiol 29, 236–243, doi:10.1152/jappl.1970.29.2.236 (1970).

56 Brooks, S. M. & Barber, M. O. Changes in closing volume measurement after isoproterenol inhalation. Am Rev Respir Dis 109, 198–204, doi:10.1164/arrd.1974.109.2.198 (1974).

57 Crawford, A. B., Makowska, M. & Engel, L. A. Effect of bronchomotor tone on static mechanical properties of lung and ventilation distribution. J Appl Physiol (1985) 63, 2278–2285, doi:10.1152/jappl.1987.63.6.2278 (1987).

58 Gazzola, M. et al. Airway smooth muscle tone curbs hyperresponsiveness in experimental asthma. BioRxiv, 1–22, doi:10.1101/2024.07.05.602208 (2024).

59 Bossé, Y., Chin, L. Y., Paré, P. D. & Seow, C. Y. Adaptation of airway smooth muscle to basal tone: relevance to airway hyperresponsiveness. Am J Respir Cell Mol Biol 40, 13–18, doi:10.1165/rcmb.2008-0150OC (2009).

